# Amount of antigen, T follicular helper cells and quality of seeder cells shape the diversity of germinal center B cells

**DOI:** 10.1101/2022.10.26.513835

**Authors:** Amar K. Garg, Tanmay Mitra, Marta Schips, Arnab Bandyopadhyay, Michael Meyer-Hermann

**Affiliations:** Department of Systems Immunology and Braunschweig Integrated Centre of Systems Biology, Helmholtz Centre for Infection Research, Braunschweig, Germany; Institute for Biochemistry, Biotechnology and Bioinformatics, Technische Universität Braunschweig, Braunschweig, Germany

**Author notes:** Correspondence: Tanmay Mitra, Michael Meyer-Hermann. These authors contributed equally to this work (alphabetical order on the basis of surnames) (Tanmay Mitra, Marta Schips).

**Keywords:** germinal center, clonal diversity, clonal dominance, founder cells, T follicular cells, agent-based model, mathematical modeling

## Abstract

A variety of B cell clones seed the germinal centers, where a selection stringency expands the fitter clones to generate higher affinity antibodies. However, recent experiments suggest that germinal centers often retain a diverse set of B cell clones with a range of affinities and concurrently carry out affinity maturation. Amid a tendency to flourish germinal centers with fitter clones, how several B cell clones with differing affinities can be concurrently selected remains poorly understood. Such a permissive selection may allow non-immunodominant clones, which are often rare and of low-affinity, to somatically hypermutate and result in a broad and diverse B cell response. How the constituent elements of germinal centers, their quantity and kinetics may modulate diversity of B cells, has not been addressed well. By implementing a state-of-the-art agent-based model of germinal center, here, we study how these factors impact temporal evolution of B cell clonal diversity and its underlying balance with affinity maturation. While we find that the extent of selection stringency dictates clonal dominance, limited antigen availability on follicular dendritic cells is shown to expedite the loss of diversity of B cells as germinal centers mature. Intriguingly, the emergence of a diverse set of germinal center B cells depends on high affinity founder cells. Our analysis also reveals a substantial number of T follicular helper cells to be essential in balancing affinity maturation with clonal diversity, as a low number of T follicular helper cells impedes affinity maturation and also contracts the scope for a diverse B cell response. Our results have implications for eliciting antibody responses to non-immunodominant specificities of the pathogens by controlling the regulators of the germinal center reaction, thereby pivoting a way for vaccine development to generate broadly protective antibodies.

## 1 INTRODUCTION

Hundreds of B cell clones seed the germinal center (GC) in B cell follicles of draining lymph nodes after infection or immunization (1, 2).A selection stringency mediated by the competition among B cell clones and their somatic mutants within a GC results in expansion of the fitter clones (3, 4). This generates higher affinity antibodies through repeated rounds of somatic hypermutation (SHM) of B cell receptors (BCRs) in the dark zone (DZ) and positive selection in the light zone (LZ) (5, 6, 7). LZ B cells (centrocytes), based on their BCR affinity, capture antigen from follicular dendritic cells (FDCs) and display internalized antigen as peptide-major histocompatibility complex class II (pMHCII) complexes to a limited number of T follicular helper (Tfh) cells (8) to get refuelled (9, 10). Following positive selection in the LZ, successful B cells re-migrate to the DZ for further rounds of proliferation and SHM (4, 5, 11, 12, 13). If GCs operated in a purely deterministic fashion, B cell clones with the highest affinity would eventually outcompete and monopolize the GC with its clonal burst sweeping away all other clones (14). However, an exceedingly affinity-stringent selection like this may target superficial non-neutralizing epitopes and fail to elicit a neutralizing immune response (15). Thus, retaining a diverse pool of B cell clones in GCs without compromising affinity maturation could be crucial in generating a broad response (13, 16, 4). In recent experiments, different GCs arising from identical conditions even within the same lymph node showed different degrees of clonal diversity loss as GC reactions progressed (2). While some GCs remained permissive for a significantly varying number of co-existing B cell clones with a range of affinities (2, 17), some were more homogeneous (2). Stochastic effects among competitors and resources may contribute to the observed diversity of GC responses (18, 19). However, amid a tendency to flourish GCs with fitter clones, how the selection of non-optimal B cell clones with differing affinities can be tuned remains poorly understood. Such a permissive selection may allow precursor clones targeting non-immunodominant epitopes, which are often rare and of low-affinity (20, 21), to somatically hypermutate and increase the diversity of B cell responses (17).

Immunodominance hinders the generation of a diverse set of neutralizing antibodies (nAbs) and that of broadly nAbs (bnAbs) proficient in eliciting a response to the evolutionary conserved epitopes (22). In the absence of concurrent recognition of several epitopes, pathogens may escape neutralizing responses as observed for influenza, HIV and SARS-COV-2 (23, 24, 25). Immunization studies retaining a greater number of unique B cell lineages in GCs, such as in the case of slow delivery immunization, were shown to target a broader range of epitopes (26) and were suggestive of improved neutralization involving a polyclonal antibody response (24). Thus, a clonally diverse GC B cell response is critical for eliciting immune responses to variants of a refractory pathogen and development of cross-strain vaccines. Although natural emergence of bnAbs was observed rarely in a fraction of chronically infected populations such as in 10% - 30% HIV+ patients (27, 28), efforts to evoke bnAbs through immunization have been of limited success (29). However, in a recent study, increasing the quantity of HIV-1 envelope glycoprotein (Env)-specific CD4 T cell help resulted in better recruitment and response of rare bnAb precursor B cells following Env-trimer immunization (30). In non-human primates, slow delivery immunization protocols with HIV-Env protein could develop potent nAbs targeting a wide range of epitopes following enhancement of Tfh cells and GC B cell responses (26). Retention of low-affinity B cell clones in GCs may also contribute to clonal diversity by providing them ample time to get refuelled by Tfh cells (10).

The aforementioned results are reminiscent of the importance of the dynamic interplay among the constituent elements of GCs, their temporal kinetics and quantity in modulating the breadth and diversity of B cell response. By implementing a state-of-the-art agent-based model of the GC (see Methods), we studied how the GC components and their intertwined dynamics would impact temporal evolution of B cell clonal diversity and its underlying balance with affinity maturation. While the extent of selection stringency dictated clonal dominance, intriguingly, maintenance of clonal diversity depended on a critical amount of available antigen on FDCs, GC founder cells of high-affinity and a substantial number of Tfh cells. Constraints therein resulted in a greater loss of clonal diversity with GC evolution. Our results have implications in eliciting GC antibody responses to non-immunodominant epitopes of the pathogens by tuning the regulators of the GC reaction, thereby pivoting a way for vaccine development to generate broadly protective antibodies.

## 2 MATERIALS AND METHODS

We developed an agent-based simulation of the GC that allows for the analysis of GC B cell diversity in a primary response. A brief overview of the simulation framework is provided here with the details presented in the Supplementary Material. The simulation starts with the generation of a 3-dimensional space on which different motile cell objects (FDCs, Tfh cells and B cells) are placed. The sequences of the B cell receptors and antigens are defined using an abstract shape space (31). The mutation distance between these sequences is used to calculate binding probability or the affinity of the B cell for the antigen. GC clonal composition is simulated by allowing the entry of unique GC B cell clones as founder cells at a rate of 2 cells per hour till 96 hours resulting in ~ 180 - 200 founder B cells (19). These founder B cells are assumed to be clonally distinct. The number of founder cells is in accordance with experiments (2). The clonal identity of B cells is maintained regardless of mutations and passed on to their daughter cells. We assume founder cell affinities and numbers to be independent of all other GC parameters. Upon entry into the GC, B cells proliferate with mutations. They then attempt to acquire antigen on the FDCs in proportion to their binding probabilities. Bound antigen is removed from the FDC, thus, reducing antigen availability over time. Further, antibodies derived from GC B cells that differentiated to plasma cells bind antigen on FDCs and make it less accessible to B cells, a phenomenon known as endogenous antibody feedback (32). While the results reported here are obtained with endogenous antibody feedback as it exists naturally in GC, simulations in the absence of it are found to be qualitatively similar (for an example, see Supplementary Material, Fig. S2).

The antigen is assumed to consist of a single epitope and to remain unmutated over the time of the GC reaction. We alter various GC handles which are relevant to GC diversity. To calculate GC diversity over time, we track the populations of different B cell clones through the course of the GC reaction and calculate the founder cell Shannon Entropy (fcSE) at time t as:

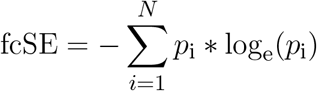

where *N* is the total number of distinct B cell clones in a GC at time *t* and *p*_i_ is the total number of cells of the *i*^th^ clone (*χ*_i_) divided by the total number of B cells (*χ*_total_) at time t:

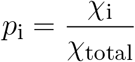

In addition, we also calculate the clonal dominance by taking the frequency distribution of the dominant B cell clone’s mole fraction in independent GC simulations. Finally, we also calculate the cumulative GC Response (CGR) as a product of average GC B cell affinity and fcSE. This additional metric allows analysis of the GC response incorporating both affinity and diversity.

## 3 RESULTS

### 3.1 Limited antigen availability expedites loss of GC diversity

Affinity dependent antigen acquisition by B cells is crucial for their survival and fate in the GC (33). Here, we investigated how antigen availability on FDCs may impact GC evolution and retention of a diverse pool of GC B cells with different affinities. We analysed three scenarios with low (1000), intermediate (3000) and high (5000) units of antigen per FDC.

The process of affinity maturation was similar (Fig. 1A) across these three settings. As we increased the initial antigen amount, the maximal size of GCs was enhanced and the plateau after the peak was extended (Fig. 1B). Additionally, the GCs started to collapse earlier with less antigen due to limited amount of free antigen on FDCs (see Supplementary Material, Fig. S1A). The number of output cells was also larger for larger antigen amounts (see Supplementary Material, Fig. S1B).

**Figure 1.**
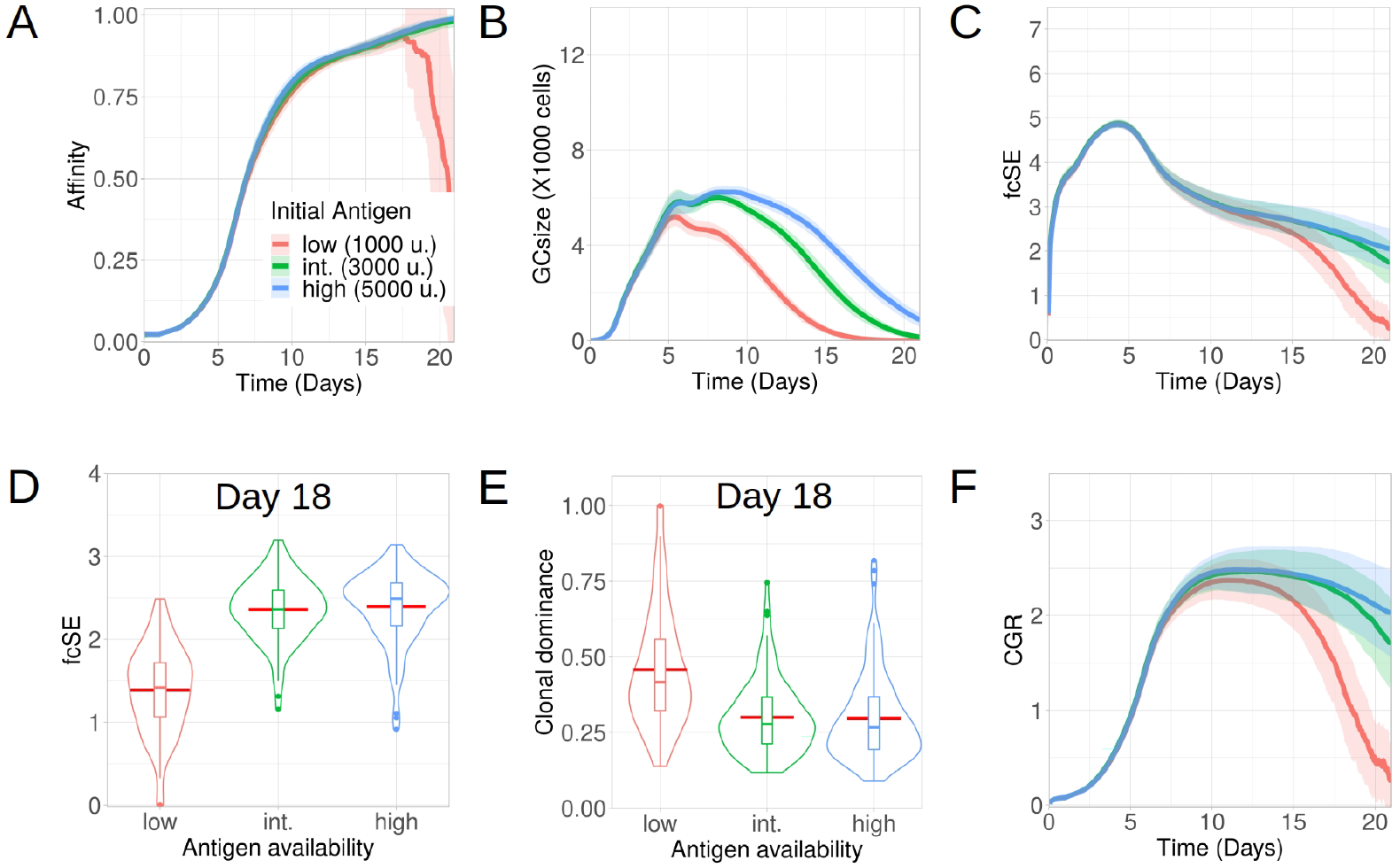
Lower antigen expedites loss of clonal diversity in GCs. Simulations were done with changing the initial antigen amount on FDC as low (1000 units in red), intermediate (3000 units in green) and high (5000 units in blue): (A) affinity of GC B cells, (B) GC size, (C) founder cell Shannon Entropy (fcSE), (D) violin plot of fcSE at day 18, (E) violin plot of clonal dominance at day 18 and (F) cumulative GC response (CGR). Mean (continuous lines) and standard deviation (shaded area) of simulations for a total of 100 simulated GCs are shown. The box plots of the violin plots show the median (horizontal line inside the box), 25 and 75 percentiles, the mean (horizontal red line) and the outlier points as dots. The relevant system parameters can be found in Table. S1 of the Supplementary Material.

The fcSE which we measured using founder cell clones to depict GC B cell diversity increased in a similar fashion at the expansion phase (till ~day 5) (Fig. 1C). Afterwards, the GC diversity waned gradually for all cases as the GC matured (Fig. 1C) because of clonal competition leading to selection of fitter clones. This was similar to experimental observations (12) where early GCs, in contrast to late GCs, were more enriched with diverse B cell clones. However, the rate of this loss in clonal diversity was slower for higher antigen amounts. Notably, following the selection phase (till ~ day 12), the fcSE contracted significantly for the *in silico* study with low amount of antigen due to the loss of B cell clones leading to a drop in the GC size.

Individual GC trajectories of the fcSE (see Supplementary Material, Fig. S1E–G) showed that many GCs in the case of low antigen go to zero fcSE indicating the transition from a highly polyclonal GC composition to a monoclonal composition. In contrast, some degree of polyclonality was maintained for the case of high antigen during the period of 21 days of GC reaction. Correspondingly, the violin plot of the fcSE values at day 18 (Fig. 1D) depicted a lower median value for the case of low antigen as compared to the other scenarios. The violin plot for clonal dominance (fraction of B cells stemming from the most dominant clone) at day 18 (Fig. 1E) revealed the clonal dominance to be higher for the case of low antigen as would be expected from the fcSE values. The CGR (Fig. 1G) peaked between day 10 and day 12 before declining at differing rates inversely correlated with the initial antigen amount. Hence, our results suggested that increasing the initial antigen availability on FDCs could prolong the GC lifetime and slow down the loss of GC B cell diversity.

### 3.2 GC diversity is enhanced by high affinity founder cells

In addition to the amount of antigen presented by the FDCs, the initial affinity of the BCRs of GC founder cells determines the fate of the GC reaction (34, 35, 36, 37, 38).

Although entry of B cells into a GC is affinity dependent (39), the onset of a GC reaction is often promiscuous in nature, permitting the participation of low-affinity B cells (40, 41, 42). Here, we investigated how tweaking the affinity of GC founder cells altered the evolution of clonal diversity, affinity maturation and CGR during the GC reaction. To do so, we increased the mutation distance (i.e., decreased affinity) of the founder cells in the shape space (see Methods) from the default distance between 5 and 6 (high affinity) to a mutation distance between 6 and 7, or 7 and 8 (intermediate and low affinity, respectively) (Fig. 2). Affinity maturation for the case of low affinity founder cells lagged behind and GC B cells had lower affinity throughout the course of the GC reaction.

**Figure 2.**
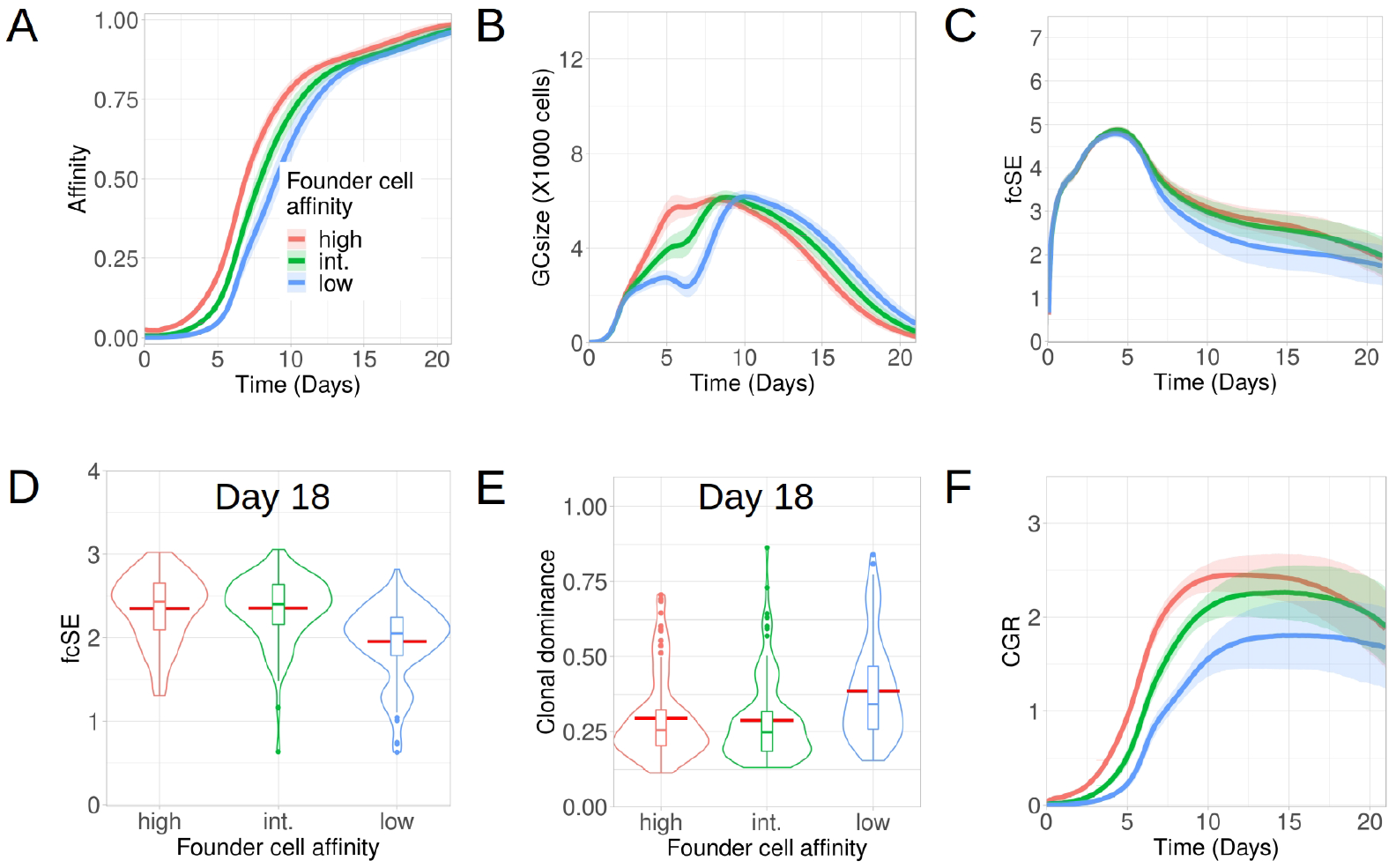
Low affinity founder cells expedite diversity loss within GCs. Simulations were done with changing the affinity of founder cells as high (mutation distance 5 to 6 in red), intermediate (mutation distance 6 to 7 in green) and low (mutation distance 7 to 8 in blue): (A) affinity of GC B cells, (B) GC size, (C) founder cell Shannon Entropy (fcSE), (D) violin plot of fcSE at day 18, (E) violin plot of clonal dominance at day 18 and (F) cumulative GC response (CGR). Mean (continuous lines) and standard deviation (shaded area) of simulations for a total of 100 simulated GCs are shown. The box plots of the violin plots show the median (horizontal line inside the box), 25 and 75 percentiles, the mean (horizontal red line) and the outlier points as dots. The relevant system parameters can be found in Table. S1 of the Supplementary Material.

The corresponding GCs faced constriction in their size early around day 6 (Fig. 2B) as a consequence of stronger selection stringency emerging from a limited amount of antigen captured from FDCs and restricted Tfh help, owing to their decreased affinity. Such a stringent clonal competition wiped out some of the GC B cell clones, thereby leading to a shrunken GC size early and an overall loss of B cell diversity as depicted by the fcSE plot (Fig. 2C).

Correspondingly, the violin plot of the fcSE values at day 18 (Fig. 2D) showed a lower median value of the overall spread with low affinity founder cells. In agreement, the clonal dominance at day 18 (Fig. 2E) showed a higher median value of the overall spread with low affinity founder cells. Finally, the CGR was curtailed when GCs were seeded by lower-affinity founder cells (Fig. 2F). Our results, thus, insinuated that the retention of a diverse set of B cells with varying affinities in GCs was promoted by high affinity GC founder cells.

### 3.3 High affinity external antibodies reduce GC diversity for lone epitopes

Specific soluble antibodies of endogenous and exogenous origin may “mask” their corresponding epitopes and/or directly compete in GCs with B cells having BCRs of similar specificities (32).

While selective masking of a dominant epitope by its specific antibody provides a possibility to promote affinity maturation of a second less-accessible epitope (43, 44), such antibody mediated feedback can increase the selection stringency of B cells and accelerate the emergence of fitter clones by reducing its probability of antigen acquisition as shown in the context of anti-hapten response (32, 45). The impact of external antibody injection on GC B cell diversity in response to a single epitope is rather unexplored.

Apart from analysing the scenario without any exogenous antibody feedback (the null case), we studied how external injection of low (*K*_D_ = 500 nM) and high affinity (*K*_D_ = 6 nM) antibodies at the start of the GC reaction affect the GC response and its B cell diversity *in silico*. In accordance with experimental observations (32), we found that injection of high affinity antibodies increased the selection pressure and resulted in an effectively faster affinity maturation having fitter clones despite showing an initial delay in increasing the average affinity of the GC B cells (Fig. 3A). In addition, such a strict selection bias for the high-affinity B cells led to the extinction of low affinity B cell lineages and constricted the GC size early during the GC reaction (Fig. 3B). Loss of low affinity B cell clones reduced the clonal diversity. For the scenario with injection of high affinity antibodies, this is reflected in the fcSE (Fig. 3C,D), with a substantially reduced mean fcSE, as well as in the increased clonal dominance (Fig. 3E). Consequently, the CGR was found to be substantially lower for the case of exogenous high affinity antibody feedback (Fig. 3F).

**Figure 3.**
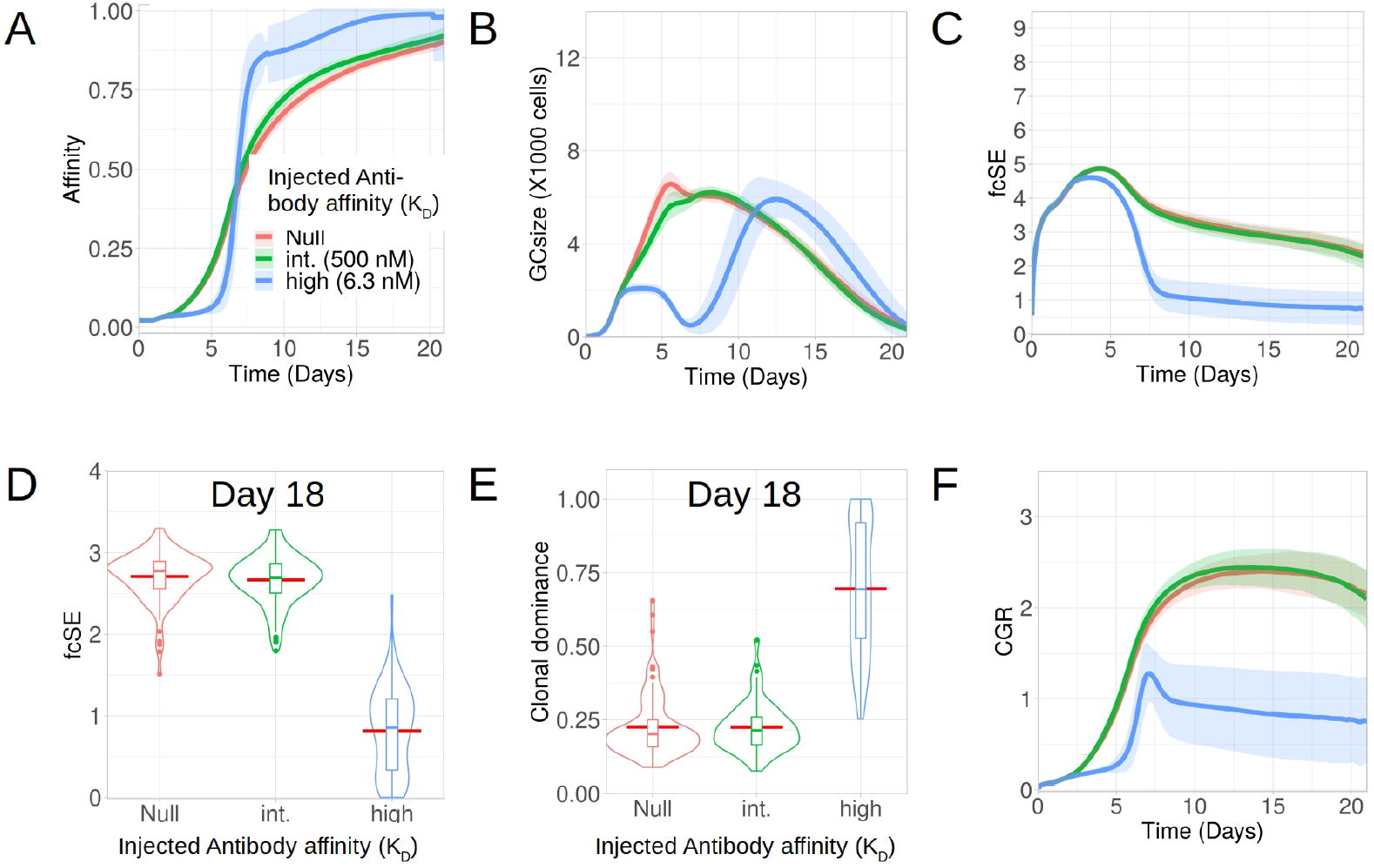
Exogenous high affinity antibody feedback reduces GC diversity. Simulations were done with changing the affinity of exogenous antibody injection provided at the zeroth day of GC evolution as null (no injection in red), intermediate (500 nM in green) and high (6.3 nM in blue): (A) affinity of GC B cells, (B) GC size, (C) founder cell Shannon Entropy (fcSE), (D) violin plot of fcSE at day 18, (E) violin plot of clonal dominance at day 18 and (F) cumulative GC response (CGR). Mean (continuous lines) and standard deviation (shaded area) of simulations for a total of 100 simulated GCs are shown. The box plots of the violin plots show the median (horizontal line inside the box), 25 and 75 percentiles, the mean (horizontal red line) and the outlier points as dots. The relevant system parameters can be found in Table. S1 of the Supplementary Material.

We also simulated GCs with external antibody injection at day 6 (see Supplementary Material, Fig. S3). Similar to the case of early antibody injection, the delayed antibody injection also resulted in the hallmarks of increased selection stringency (faster affinity maturation, constriction of GC size after antibody injection and reduced diversity), although to a lesser extent. This is due to the injection being performed when affinity maturation in the GC already progressed, such that the GC B cells are more competitive (32)). Taken together, our results demonstrated in the case of B cell response to a single epitope that injection of high affinity antibody induced selection stringency accelerating the process of affinity maturation, and reduces GC B cell diversity by early elimination of low affinity clones.

### 3.4 Limited Tfh cell numbers stunt affinity maturation and reduce GC diversity

B cells compete for interacting with Tfh cells in the LZ to receive signals for survival and proliferation (3). Thus, the magnitude of Tfh help is a limiting factor in the GC reaction mediating the selection of the B cells (46, 47, 48). Therefore, an alteration in the number of Tfh cells may regulate the selection stringency and can have implications in maintaining a diverse pool of GC B cells. Here, we analysed the evolution of GC diversity for three *in silico* scenarios having differing number of Tfh cells, viz., with 300 (high), 200 (intermediate, this was the default number of Tfh cells in the results discussed so far) and 100 (low). While affinity maturation for the high and intermediate number of Tfhs was identical, the process was significantly slower and less efficient in attaining higher average affinity of the surviving GC B cells for the low number of Tfh cells (Fig. 4A). Overall, GC sizes and their peaks were also reduced with less Tfh cells (Fig. 4B).

**Figure 4.**
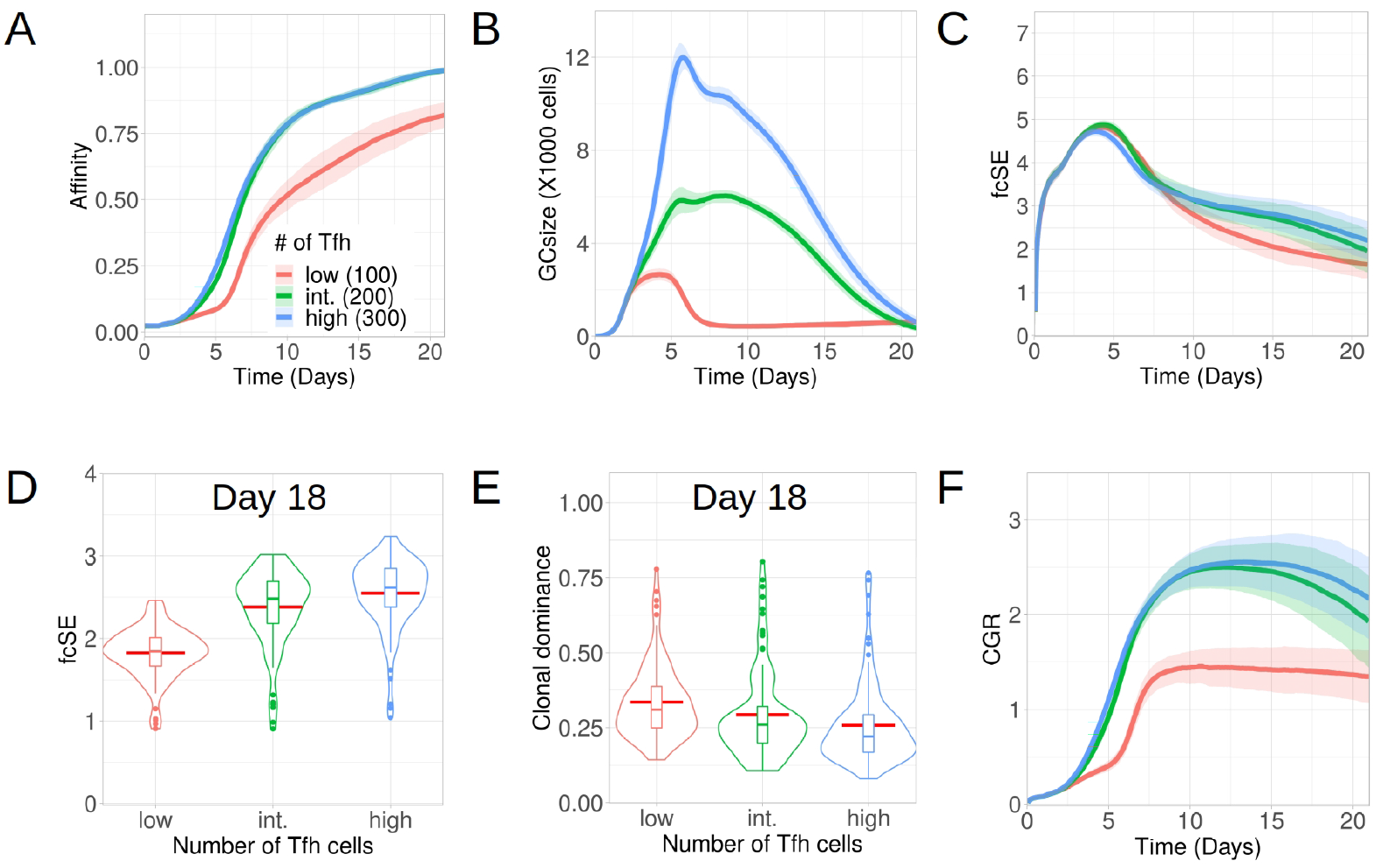
Low Tfh cell numbers stunt affinity maturation and increase loss of GC diversity. Simulations were done with changing Tfh numbers as low (100 units in red), intermediate or default (200 units in green) and high (300 units in blue): (A) affinity of GC B cells, (B) GC size, (C) founder cell Shannon Entropy (fcSE), (D) violin plot of fcSE at day 18, (E) violin plot of clonal dominance at day 18 and (F) cumulative GC response (CGR). Mean (continuous lines) and standard deviation (shaded area) of simulations for a total of 100 simulated GCs are shown. The box plots of the violin plots show the median (horizontal line inside the box), 25 and 75 percentiles, the mean (horizontal red line) and the outlier points as dots. The relevant system parameters can be found in Table. S1 of the Supplementary Material.

One might have expected that with low Tfh cell numbers competition is increased and, thus, affinity maturation would be accelerated. This tendency was counter-acted by the rather strong effect on the overall size of the GC response (Fig. 4B), which did not allow for the number of surviving B cells required to evolve high affinity B cells(see Supplementary Material, Fig. S4).

Because of early constriction in GC size, the fcSE dropped post ~ day 7 for low Tfh cell numbers (Fig. 4C).The violin plots of fcSE (Fig. 4D) and clonal dominance (Fig. 4E) at day 18 showed opposing trends with increasing number of Tfh cells, indicating more diverse GCs for higher Tfh cell counts. However, the differences in these metrics were rather insignificant between intermediate and high numbers of Tfh cells. Consequently, the CGR (Fig. 4F) was also lower for low Tfh cell counts.

We also investigated whether the number of divisions of B cells underwent any change due to alteration in the quantity of Tfh cells. The normalized frequencies of B cell divisions (see Supplementary Material, Fig. S6A–C) and the mean number of B cell divisions (see Supplementary Material, Fig. S6D) were similar in all cases. Notably, the mean number of divisions was between 2 and 2.5, which was in agreement with earlier experimental observations (12).

The simulations were repeated for a wider range of Tfh cell counts and the corresponding fcSE and CGR were reported at day 18 (see Supplementary Material, Fig. S5A–B). While both these metrics were lower for lower numbers of Tfhs (100 and 150 Tfhs), a saturation in fcSE and CGR was observed as we increased the number of Tfh cells indicating the absence of improved GC response beyond a threshold number of Tfh cells. Thus, our results suggested that a limited number of Tfh cells could negatively affect affinity maturation, constrict the GC size and reduce GC diversity.

### 3.5 Low quality Tfh cells stunt GC response and accelerate loss of diversity

Next, we investigated how altering the quality of the Tfh cells would impact on the GC response and GC B cell diversity. B cells compete to acquire critical signals from Tfh cells to get positively selected in GCs. To invoke changes in Tfh quality in our *in silico* framework, the amount of signals acquired by a B cell following the interaction with a Tfh cell was multiplied with a signal multiplier value. Low quality Tfh cells have a lower expression of co-stimulatory signalling factors and thus, B cells would require a larger number of interactions and overall longer duration of Tfh signalling to get selected. We implemented this by lowering the value of the Tfh signal multiplier. In contrast, a higher value of Tfh signal multiplier would allow for B cell selection with a lower overall duration of Tfh signalling.

Accordingly, we studied the GC dynamics for both low and high quality Tfh cells with low (0.6 in red), intermediate low (0.8 in green), default (1 in cyan) and high (1.2 in purple) Tfh signal multiplier values (Fig. 5). Affinity maturation was observed to be relatively robust and declined only for the lowest value of Tfh signal multiplier (Fig. 5A). In contrast, the GC size showed strong dependence on the Tfh signal multiplier value and progressively increased with higher quality Tfh cells (Fig. 5B). The fcSE and clonal dominance declined significantly only for the lower value of Tfh signal multiplier (Fig. 5C, D and E) and were otherwise robust. CGR was lower for low quality Tfh cells (Fig. 5F). The normalized frequencies of B cell divisions were found to be similar in all cases and the mean number of B cell divisions was between 2 and 2.5 as before (see Supplementary Material, Fig. S7). Taken together, our results showed that below a certain quality of Tfh cells, the GC response falters showing slower and inefficient affinity maturation, reduced GC size, accelerated loss of B cell diversity and a reduced CGR. Increase in Tfh cell quality beyond this threshold shows a relatively robust GC response.

**Figure 5.**
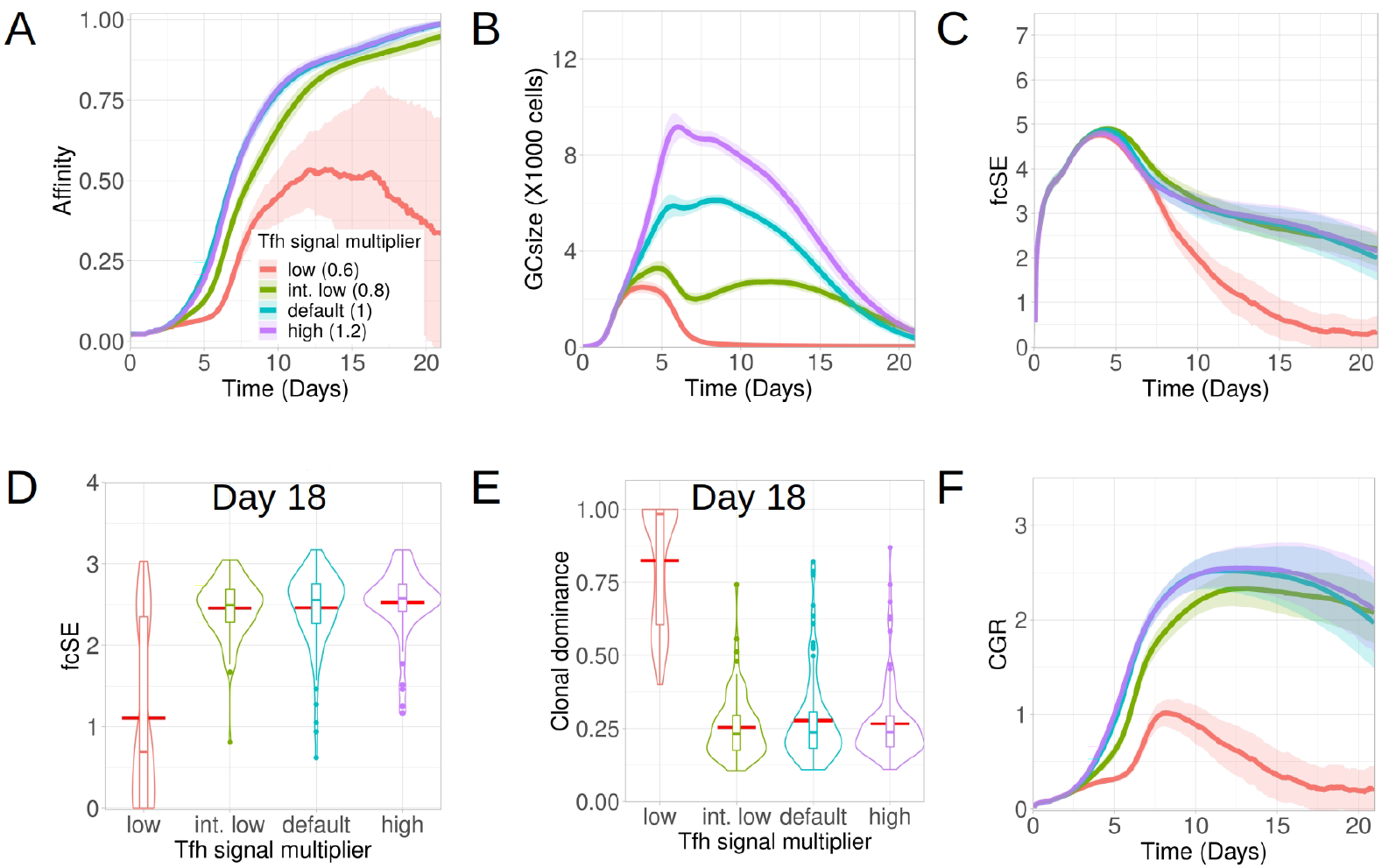
Low Tfh cell quality stunts affinity maturation and increases loss of GC diversity. Simulations were done with changing the Tfh signal multiplier as low (0.6 in red), intermediate low (0.8 in green), default (1 in cyan) and high (1.2 in purple): (A) affinity of GC B cells, (B) GC size, (C) founder cell Shannon Entropy (fcSE), (D) violin plot of fcSE at day 18, (E) violin plot of clonal dominance at day 18 and (F) cumulative GC response (CGR). Mean (continuous lines) and standard deviation (shaded area) of simulations for a total of 100 simulated GCs are shown. The box plots of the violin plots show the median (horizontal line inside the box), 25 and 75 percentiles, the mean (horizontal red line) and the outlier points as dots. The relevant system parameters can be found in Table. S1 of the Supplementary Material.

## 4 DISCUSSION

We developed an agent-based model for the GC response to investigate how retention of a diverse array of B cells with differing affinities depends on the availability of antigen on FDCs, initial affinity of the GC founder cell pool, epitope masking by specific antibodies, and the quantity and quality of the Tfh cells. By analysing the intertwined dynamics of these critical GC components, we studied their impact on temporal evolution of B cell clonal diversity and its underlying balance with affinity maturation. We found that limited antigen availability on FDCs expedites loss of GC diversity during the GC evolution. We further showed that emergence of a diverse set of GC B cells requires the founder cells to have good BCR affinity. Additionally, in the case of B cell response to a single epitope, our results depicted that high affinity external antibody feedback reduced GC diversity through early extinction of low affinity B cell clones. By assuming a different number of Tfh cells in *silico,* we showed that a minimum number of Tfhs was needed for proper affinity maturation and retention of GC B cell diversity.

Whereas FDCs loaded with limited amount of antigen resulted in a rapid loss of B cell diversity as a consequence of accelerated GC shutdown, a surplus of antigen availability promoted retention of a diverse set of B cell clones for longer duration inside GCs, but did not lead to a significant increase in the peak clonal diversity during the GC evolution. Slow delivery immunization protocols that promoted antigen retention in the lymph nodes and resulted in sustained antigen availability inside the GC (49) were shown to enhance GC size and clonal diversity, thereby enhancing neutralizing antibody responses (49, 26). This was previously thought to arise from the elongation of a B cell recruitment time-window that provided the B cells with rare precursor frequencies a chance to be recruited in the GC (50). Dynamically augmented Tfh help was also considered to contribute to such an observation (49, 26). Although we did not explicitly model slow delivery immunization mechanisms here, we demonstrated that the decline of GC size due to antigen starvation could accelerate the loss of clonal diversity, possibly restraining permissive selection of low-affinity B cells, making it one of the possible confounding factors for emergence of fewer nAbs. Notably, we observed retention of diverse B cell clones for longer duration and an enhanced GC size without prolonging B cell recruitment time window or invoking augmented Tfh help as we provided GC with more antigen *in silico*. Sustained availability of antigen might have overcome antigen starvation and contributed to robust nAbs generation. Our results, thus, provided an alternative explanation for broader B cell responses observed in slow delivery immunization studies.

We further found, when GCs were seeded with comparatively lower-affinity founder B cells, the resulting GC response was more stringent towards selecting the fitter clones thereby sacrificing a fair bit of B cell clonal diversity. In case of higher-affinity founder cells seeding the GC, a relatively faster and efficient affinity maturation along with retention of a wide range of B cell clones with varying affinities resulted in a diverse GC and a better cumulative GC response. The aforementioned requirement for a clonally diverse GC might be related as to why it can be difficult to elicit a natural bnAb response (51), provided bnAb precursor cells can have lower affinity compared to other non-bnAb precursor cells (52).

Specific antibody induced epitope-masking (antibody feedback) (32) was previously shown to alter the focus of affinity maturation to a second but less available epitope in *silico* (43). Thus, selective masking of a dominant epitope by specific antibody is a plausible mechanism to promote affinity maturation of the less-accessible epitope, thereby creating a broader B cell response. Indeed, studies investigating a Malaria vaccine suggested, whereas the recall B cell response to immunodominant PfCSP (Plasmodium falciparum circumsporozoite protein) repeat region was inhibited by antibody feedback, subsequent boosting expanded the subdominant responses to PfCSP C-terminal regions (44). However, in the context of antibody responses to a single epitope, in *silico* injection of high affinity external antibody turned out to affect GC diversity negatively as there is no other epitope to focus the GC response to. In such a scenario, external antibody feedback induced selection stringency and by a transient constriction of GC size early during the GC response, reduced clonal diversity. Immunization with external antibodies, which is more commonly known as passive immunizations, was first used for the treatment of the 1918 influenza pandemic (53). More recently, such a treatment option was also explored for Ebola (54), Influenza (55) and the SARS-COV-2 pandemic (56, 57). Depending on how specifically an antibody can mask its corresponding epitope and the number of antigenic epitopes for which the B cell response is triggered, the outcome can be very different. During B cell responses to a single epitope, while passive immunization can speed up affinity maturation, it can suppress GC diversity. Thus, it can be potentially detrimental for the evolution of important neutralizing responses, particularly if the specific B cell lineages are compromised in the early GC reaction through external antibody feedback.

Positive selection and expansion of higher affinity B cells during affinity maturation occur at the expense of their lower affinity competitors (3, 4). The lower affinity B cell clones undergo apoptosis at a higher rate than B cells with intermediate affinities because of limited amount of antigen capture and, consequently, constricted Tfh help (58). The bottleneck of low affinity B cell survival, thus, hinges upon the affluence of resources and outcome of the relative competition among the B cells. As bnAb precursor B cells are rare and of relatively lower affinity (30, 52), their survival and SHM depend on relaxing the selection stringency in GC. Dynamically increasing the number of Tfh cells during an ongoing GC reaction might be one of the possible mechanisms that could help retaining low-affinity B cells inside GC for a longer duration by reducing selection stringency and provide them the chance to receive Tfh help, thereby contributing to clonal diversity Although we do not explicitly explore this here, our finding that a critical number of Tfh cells and certain level of antigen availability are needed to maintain clonal diversity suggests this. Our analysis revealed a substantial number of Tfh cells to be essential in balancing affinity maturation with clonal diversity, as a low number of Tfh cells impeded affinity maturation and contracted the scope for a diverse GC. Consequently, it implies that during a GC reaction when antigen is gradually taken up by the B cells depending on their BCR affinity to the pMHCs presented by FDCs, a contemporaneous expansion of Tfh cells would succour the less fit B cell clones to survive the competition. Hence, the bidirectional help between Tfh and B cells where selective expansion of the Tfh cells would depend upon signals received from the B cells may contribute to clonal diversity, a field worth exploring experimentally. As receiving Tfh help is a critical gridlock for entry of B cells into the GC and their survival therein (39, 50), such a dynamic regulation of Tfh cell quantity as previously observed in experiments (59) would seem advantageous in broadening the breadth of antibody neutralization. Indeed, evidence suggests improvement of neutralization titres when the number of Tfh cells was increased (60, 26).

In the GC settings studied here, clonal diversity is robustly reduced during the second week of the GC reaction. Provided the export of GC derived long-lived IgG1 memory B cells (MBCs) peaks prior to attaining a fully-fledged GC (61), these MBCs should consist of a diverse set of low-affinity non-immunodominant clones capable of recognizing epitope-variants (62, 63, 64). As MBCs hardly participate in subsequent GC responses to the same antigen (65), a relatively early export of clonally diverse MBCs during the primary response would support diversification of an initial pool of antibodies upon re-activation. On the contrary, long-lived plasma cells are predominantly derived later from fitter clones with more rounds of SHM (61, 66, 65, 67), consequently resulting in a less-diverse clonal pool (17). An intriguing possibility that emerges from these observations is whether GC is trying to optimize its cumulative response across different timescales while exporting both effector outputs, viz., MBCs and plasmablasts having distinct features regarding clonal diversity and affinity.

A less studied question in the context of T follicular cells, is how the presence of T follicular regulatory (Tfr) cells and their relative abundance to that of Tfh population shape clonal diversity in GC. Data from SARS-COV-2 protein vaccination studies suggest a role of Tfr cells in promoting the contribution of SARS-CoV-2-specific clones in GC by restraining competition (68). However, in chronic GCs emerging in autoimmune disorders wherein the ratio of Tfh and Tfr cells is mostly increased (69, 70, 71, 72), little is known about clonal diversity. Notably, these non-resolving GCs home distinct GC reactions seeded by B cell clones specific for different autoantigens (73), and thus, exploring these scenarios using in-vivo studies is challenging. Whether high Tfh/Tfr ratio leads to generation of a broad range of autoantibodies through epitope spreading or such an altered ratio is a consequence of the humoral response trying to restore the homeostasis, is alluring. One possible extension of our model could be to study this question designing *in silico* experiments.

While a high level of diversity in GC B cell response would be deemed advantageous for eliciting immune responses to variants of a pathogen, it may endanger an organism’s own cells by provoking an autoimmune response if the pathogenic epitopes resemble self-peptides substantially. Our simulations also illustrate the contexts attaining limited B cell diversity in GC, which might be related to understanding the checkpoints on autoimmunity. As the founder B cells initiate GC in a T-cell dependent manner, ideally, avoidance of profound T-cell autoimmunity would congruently imply the affinity of the GC founder B cells to be low in such a context. Our results suggest that the resulting GC response, in the case of low-affinity founder cells, can be skewed towards the most optimal clone with the highest affinity to the specific epitopes rather than generating a diverse array of B cells potentially capable of reacting to self-tissues. It is, thus, intriguing to consider a GC as a possible machinery that regulates its B cell diversity to meet these two opposing needs of ensuring a robust and flexible response while keeping tolerance of self.

Our results contribute to understanding how a GC response can retain diversity without compromising affinity maturation. The *in silico* framework can be used to design experimental set-ups suitable for generating broader GC B cell responses and to develop vaccine strategies to concentrate antibody responses to variants of refractory pathogens by controlling the most important regulators of the GC reactions.

## AUTHOR CONTRIBUTIONS

Conceptualization and supervision: MMH, TM;

Basic code and model development: MMH;

Coding, simulations and formal analysis: AKG, MS;

Final simulations and figures: AKG;

Investigation and discussion: TM, AKG, MS, MMH, AB;

Original draft: TM, AKG, AB;

Revision: TM, MMH;

## ACKNOWLEDGMENTS

We thank Dr. Sebastian C. Binder for reviewing the manuscript and for helpful suggestions.

## SUPPLEMENTARY MATERIAL

A detailed description of the agent-based model of a germinal center is provided in the Supplementary Material. In addition, a table (Table. S1) for the model parameters relevant for this study and supporting figures (Fig. S1–S7) can be found therein.

## DATA AVAILABILITY STATEMENT

The codes developed and data analysed for this study can be obtained from the corresponding authors upon request.

