## Supplementary Material for "Amount of antigen, T follicular helper cells and quality of seeder cells shape the diversity of germinal center B cells"

#### 1 GERMINAL CENTER SIMULATION MODEL

The model assumptions underlying the germinal center (GC) simulations are explained here and include the used parameter values. It follows the description of (Meyer-Hermann *et al.*, 2012) adapted to include novel features introduced since then in (Meyer-Hermann, 2014; Binder & Meyer-Hermann, 2016). Used acronyms are: DZ for dark zone, LZ for light zone, Tfh for T follicular helper cell, FDC for follicular dendritic cell.

##### Space representation

All reactions take place on a three-dimensional discretized space with a rectangular lattice with lattice constant of $\Delta x = 5\mu m$. Every lattice node can be occupied by a single cell only.

##### Shape space for antibodies

Antibodies are represented on a four ($d = 4$) dimensional shape space (Perelson & Oster, 1979). The shape space is restricted to a size of 10 positions per dimension, thus, only considering antibodies with a minimum affinity to the antigen. The optimal clone $\Phi^*$ is positioned in the centre of the shape space. A position on the shape space $\Phi$ is attributed to each B cell.

The 1-Norm with respect to the optimal clone $||\Phi - \Phi^*||_1 = \sum_{i=1}^{d} |\Phi_i - \Phi_i^*|$, i.e. the minimum number of mutations required to reach the optimal clone, is used as a measure for the antigen binding probability. The binding probability is calculated from the Gaussian distribution with width $\Gamma = 2.8$ (Meyer-Hermann *et al.*, 2001):

$$b(\Phi, \Phi^*) \quad = \quad \exp\left(-\frac{||\Phi - \Phi^*||_1^2}{\Gamma^2}\right) \quad . \qquad \text{(S1)}$$

##### B cell phenotypes

Three B cell phenotypes are distinguished: DZ B cells, LZ B cells, and output cells. The different phenotypes characterize the cell properties and are not meant as localization within the GC zones. DZ B cells divide, mutate and migrate. LZ B cells also migrate and undergo the different stages of the selection process. Output cells only migrate.

##### Founder cells

The model starts from 200 Tfh, 200 FDCs, 300 stromal cells, and no B cell. Tfh are randomly distributed on the lattice and occupy a single node each. Stromal cells are restricted to the DZ (see section *Chemokine distribution* for their function). FDCs are restricted to the upper half of the reaction sphere, occupy one node by their soma and have 6 dendrites of $40\mu m$ length each. The presence of dendrites is represented as a lattice-node property and, thus, visible to B cells. The dendrites are treated as transparent for B cell or Tfh migration such that they do not inhibit cell motility.

##### B cell influx rate

As B cell selection is not active during the first 3 days of the reaction (i.e. days 3-5 post immunisation), the first 3 days can be approximated as a B cell expansion phase. Clonality can be safely ignored, and it suffices to consider a single dividing cell type $B_i$, where $i$ denotes the generation number of the B cells. The dynamics of expansion are then described by

$$\begin{aligned} \frac{dB_1}{dt} &= s - pB_1 \\ \frac{dB_\mathrm{i}}{dt} &= 2pB_{\mathrm{i}-1} - pB_\mathrm{i} \quad \text{for } i > 1 \\ \frac{dB_\mathrm{GC}}{dt} &= 2pB_{\mathrm{i}_\mathrm{max}}\,, \end{aligned} \tag{S2}$$

where $s$ is the influx rate, $p$ the division rate, $i_\mathrm{max}$ the number of divisions per cell in the expansion phase, and $B_\mathrm{GC}$ the resulting number of GC B cells that participate in the GC reaction. The number of initial divisions is estimated by the maximum number of divisions observed upon anti-DEC205-OVA treatment, i.e. $i_\mathrm{max} = 6$ (Victora *et al.*, 2010; Meyer-Hermann *et al.*, 2012). As the division time $\ln(2)/p$ is shorter than the expansion phase $T_\mathrm{expand} = 3$ days, one may solve Eq. (S2) in steady state, yielding:

$$B_6 = 2B_5 = 4B_4 = 8B_3 = 16B_2 = 32B_1 = 32s/p\,. \tag{S3}$$

Thus, the relevant ODE becomes

$$\frac{dB_\mathrm{GC}}{dt} = 2pB_6 = 64s \implies B_\mathrm{GC} = 64st\,, \tag{S4}$$

i.e. a linear growth in time proportional to the constant influx during expansion. Note that in the steady state approximation the influx rate becomes independent of the division rate $p$, which is an implication of the assumption of a fixed number of divisions per founder cell $i_\mathrm{max}$. With the side condition of getting 9000 cells at day 3, $B_\mathrm{GC}(T_\mathrm{expand} = 72\mathrm{hr}) = 9000$, the influx rate is estimated to be

$$s = \frac{B_\mathrm{GC}(T_\mathrm{expand})}{64T_\mathrm{expand}} \approx 2\ \mathrm{cells/hr}\,. \tag{S5}$$

This corresponds to 192 B cells entering the GC in the first 4 days of expansion and building up the founder cell population of the GC reaction.

Motivated by this estimation, in the model we assumed that B cells enter the GC reaction with a probability corresponding to a rate of 2 cells per hour. New B cells are randomly positioned on the lattice (exclusively on free nodes).

The shape space position of each new B cell is randomly picked from a set of 100 shape space positions at a distance of 5 or 6 mutations to the nearest epitope.

#### Antigen-presentation by FDCs

Each FDC is loaded with $3500$ antigen portions distributed onto the lattice-nodes occupied by FDC-soma or FDC-dendrite. One antigen portion corresponds to the number of antigen molecules taken up by a B cell upon successful contact with an FDC.

#### Antigen-antibody interaction on FDCs

Possible sources of antibodies are output cells of the GC reaction and injections (see section *Antibody sources* below). Antibodies are represented in the 4-dimensional shape space with 10 positions in each direction. The quantity of interest is the amount of free antigen-epitope at each FDC site when antibodies are present and changing over time. As it is not feasible to calculate the amount of free epitope at each site for all 10,000 possible antibody types, 11 affinity bins $B_\mathrm{i}$ with $i \in [0, \ldots, 10]$ were introduced, where the affinity is defined relative to the antigen-epitope. We assumed a constant on-rate $k_\mathrm{on} = 10^6/(\mathrm{Mol\,sec})$ (Batista & Neuberger, 1998) and a variable off-rate

$$k_{\mathrm{off,i}} \;=\; \frac{k_\mathrm{on}}{10^{5.5+0.4i}} \quad , \tag{S6}$$

mimicking a dissociation constant that varies over 4 orders of magnitude. At each FDC site $x$, the chemical kinetics equation for the immune complexes $C_\mathrm{i}(x)$ formed between antibodies in bin $i$ the epitope

$$\frac{dC_\mathrm{i}(x)}{dt} \;=\; k_\mathrm{on} E(x) B_\mathrm{i} - k_{\mathrm{i,off}} C_\mathrm{i}(x) \tag{S7}$$

was solved in order to determine the amount of free epitope $E(x)$ at this site. Only this amount of epitope is available for B cells to bind antigen with probability according to Eq. (S1). Additional competitive binding probability as used in (Zhang *et al.*, 2013) was assumed here. To do this the width (Γ) for the binding probability Gaussian function in Eq. (S1) was recalculated as $\Gamma_{new}$ = Γ(1-A) with A being the average affinity of all produced (or injected) antibodies with affinities as calculated with the same Gaussian in the noncompetition limit, i.e., with width as Γ.

#### DZ B cell division

The average cell cycle duration of 7 hours of DZ B cells is varied for each B cell according to a Gaussian distribution. This is needed to get desynchronization of B cell division. The cell cycle is decomposed into four phases (G1, S, G2, M) in order to localize mitotic events if this is needed.

Each founder B cell divides a number of times before differentiating to the LZ phenotype for the first time. Six divisions was the number of divisions found in response to the extreme stimulus with anti-DEC205-OVA (Victora *et al.*, 2010; Meyer-Hermann *et al.*, 2012). Each selected B cell divides a number of times determined by the interaction with Tfh (see below, LZ B cell selection). The parameters of the interaction with Tfh are tuned such that the mean number of divisions is in the range of two (Gitlin *et al.*, 2014). This value is required in order to maintain a DZ to LZ ratio in the range of two (Victora *et al.*, 2010; Meyer-Hermann *et al.*, 2012).

A division requires free space on one of the Moore neighbors of the dividing cell. Otherwise the division is postponed until a free Moore neighbor is available.

At every division the encoded antibody can mutate with a probability of $0.5$ (Berek & Milstein, 1987; Nossal, 1992). This corresponds to a shift in the shape space to a von Neumann neighbor in a random direction. Upon selection by Tfh the mutation probability is individually reduced from $m_\mathrm{max} = 0.5$ down to $m_\mathrm{min} = 0$ in an affinity-dependent way following

$$m(b) \;=\; m_\mathrm{max} - (m_\mathrm{max} - m_\mathrm{min})\, b \;=\; \frac{1-b}{2} \tag{S8}$$

with $b$ from Eq. (S1) (Toellner *et al.*, 2002). Thus, after recycling DZ B cells can acquire reduced mutation probabilities. This mechanism is motivated by the observation that B cell receptor internalization enhances the activation of the kinase AKT (Chaturvedi *et al.*, 2011) which, in turn, suppresses activation induced cytosine deaminase (AID) (Omori *et al.*, 2006). AID is required for somatic hypermutation, such that this provides an affinity-dependent down-regulation of the mutation frequency (Dustin & Meyer-Hermann, 2012). However, there is no formal proof of this mechanism.

B cell division of B cells that previously acquired antigen and have been selected by Tfh distribute the retained antigen asymmetrically to the daughters (Thaunat *et al.*, 2012). The model assumes asymmetric division in $72\%$ of the cases, which is supported by experimental observations (see (Thaunat *et al.*, 2012) and Supplementary Figure S1 in (Meyer-Hermann *et al.*, 2012)). If division is asymmetric, one daughter gets all the retained antigen while the other gets none, which approximates the value of $88\%$ found in (Thaunat *et al.*, 2012). Mutation is suppressed in cells retaining antigen.

After the required number of divisions the B cell differentiates with a rate of in $1/6$ minutes to the LZ phenotype. All B cells that kept the antigen up to this time, differentiate to output cells, up-regulate CXCR4, and leave the GC in direction of the T zone.

#### LZ B cell selection

LZ B cells can be in the states *unselected, FDC-contact, FDC-selected, Tfh-contact, selected, apoptotic*.

##### Unselected

LZ B cells migrate and search for contact with FDCs loaded with antigen in order to collect antigen for 0.7 hours. If an FDC soma or dendrite is present at the position of the B cell, the B cell attempts to establish contact to the epitope. The binding of B cell to epitope is affinity dependent and happens with the probability $b$ in Eq. (S1). If the available number of antigen portions at the specific FDC site drops below $20$ the binding probability $b$ is linearly reduced with the number of available portions. If successful, the B cell switches to the state *FDC-contact*; otherwise the B cell continues to migrate. Further binding-attempts are prohibited for $1.2$ minutes. At the end of the antigen collection period, B cells switch to the state *FDC-selected*. If a LZ B cell fails to collect any antigen at this time it switches to the state *apoptotic*.

##### FDC-contact

LZ B cells remain immobile (bound) for $3$ minutes (Schwickert *et al.*, 2007) and then return to the state *unselected*. The counter for the number of successful antigen uptake events is increased by one and the FDC reduces its locally available antigen portions by one.

##### FDC-selected

B cells search for contact with Tfh. If they meet a Tfh they switch to the state *Tfh-contact*.

##### Tfh-contact

LZ B cells remain immobile for $6$ minutes. In this time the bound Tfh, which may also be bound to other B cells, polarizes to the bound B cell with highest number of successful antigen uptakes. Only this B cell receives Tfh signals and accumulates those. After the binding time, the B cell detaches and returns to the state *FDC-selected*. It continues to search and bind Tfh cells until the Tfh search time of 3 hours is over. Then, it switches to the state *apoptotic* if the accumulated Tfh-signalling time remained below the threshold Tfh signal which has a default value of $30$ minutes. Otherwise it switches to the state *selected*.

##### Selected

LZ B cells keep the LZ phenotype for six hours and desensitize for CXCL13, thus, perform a random walk. During that time they re-enter cell cycle and progress through the cell cycle phases. Then they recycle back to the DZ phenotype with a rate of $1/6$ minutes and memorize the amount of collected antigen, the amount of Tfh signal acquired and the cell cycle phase they have achieved by this time.

The number of divisions $P(A)$ the recycled B cells will do is derived from the amount of acquired Tfh signal $A$, which reflects the the amount of antigen collected and the affinity of the B cell receptor for the antigen, as follows:

$$P(A) = P_{\min} + (P_{\max} - P_{\min}) \frac{A^{n_P}}{A^{n_P} + K_P^{n_P}} . \tag{S9}$$

The more antigen was collected by the B cell the more divisions are induced. We set the minimum number of division to one ($P_{\min} = 1$) in order to avoid recycling events without further division. It is limited by six divisions in the best case, which is motivated by anti-DEC205-OVA experiments in which DEC205$^{+/+}$ B cells received abundant antigen which increased pMHC presentation to a maximum (Victora *et al.*, 2010). The population dynamics in vivo and *in silico* only matched when the number of divisions was increased to six in the simulation (Meyer-Hermann *et al.*, 2012) suggesting that the strongest possible Tfh signalling induces six divisions ($P_{\max} = 6$). The Hill-coefficient was set to $n_P = 1.8$.

The half value $K_P$ remained to be determined, which denotes the amount of Tfh signal acquired by B cells at which the number of divisions becomes half maximal. The amount of Tfh signal acquired varies between zero and a maximum determined by the duration of the Tfh signal collection phase, the duration of each B cell interaction with Tfh cells, and the motility of B cells and Tfh cells. The frequency distribution of Tfh signals acquired by selected and apoptotic B cells as observed in the simulations served as estimate of $A_{\max}$. B cells acquire Tfh signal between 0 and ∼1 hr. For an intermediate acquired Tfh signal of $A_0 = 0.6\ hrs$, the resulting number of divisions has to be $P_0 = 2$ in order to be in agreement with the mean number of divisions in the range of two (Gitlin *et al.*, 2014), which leads to the condition:

$$K_P \approx A_0 \left( \frac{P_{\max} - P_{\min}}{P_0 - P_{\min}} - 1 \right)^{1/n_P} \approx 1.3 \quad . \tag{S10}$$

##### Apoptotic

LZ B cells remain on the lattice for $6$ hours before they are deleted. They continue to be sensitive to CXCL13 during this time.

#### Antibody sources

##### Output cells

Output cells from the GC reaction are collected and memorized together with the affinity of the encoded antibody to the epitopes. Their life time is assumed longer than the duration of the GC reaction. Output cells are attributed to the different affinity bins for each epitope and further differentiate to an antibody forming plasma cell according to a linear rate equation with a rate of $\ln(2)$ per day, i.e. with a half life of one day.

All plasma cells produce antibodies $B_i$, attributed to bin $i$ for the epitope according to

$$\frac{dB_i}{dt} = \frac{N_{GC} r_Q}{V_{blood}} Q_i - \gamma_B B_i , \tag{S11}$$

where $r_{\rm Q} = 3\,10^{-18}{\rm mol/hour}$ is the production rate per cell (Randall *et al.*, 1992) and $\gamma_{\rm B} = \ln(2)/(14{\rm days})$ is the degradation rate of antibodies. The produced antibodies are assumed to distribute over the whole organism. The sum of all $N_{\rm GC}$ GC reactions in the organism is diluted over the mouse blood volume $V_{\rm blood}$ and adds up to the total antibody concentration found in the simulated GC. The factor $N_{\rm GC}/V_{\rm blood} = 10/{\rm ml}$ was assumed. This setting implies that antibodies are homogeneously distributed over the space of the simulated GC. Also the injection of antibodies is possible.

##### Antibody injection

Injection of antibodies is represented by an additional source term $\delta\left(t - t_{\rm inject}\right) \Delta B_{\rm i}$ in Eq. (S11) at the particular time point $t_{\rm inject}$. $\Delta B_{\rm i}$ is $\Delta B_{\rm i} = \Delta B_0$ for exactly one bin $i$ and $\Delta B_{\rm i} = 0$ otherwise.

#### Chemokine distribution

Two chemokines CXCL12 and CXCL13 are considered. CXCL13 is produced by FDCs in the LZ with 10nMol per hour and FDC while CXCL12 is produced by stromal cells in the DZ with 400nMol per hour and stromal cell. As both cell types are assumed to be immobile, chemokine distributions were pre-calculated once and the resulting steady state distributions were used in all simulations.

#### Chemotaxis

DZ and LZ B cells regulate their sensitivity to CXCL13 and CXCL12, respectively. This is true in all B cell states unless stated otherwise. All B cells move with a target speed of $7.5 \mu m/min$. This leads to a slightly lower observable average speed of $\approx 6 \mu m/min$.

B cells have a polarity vector that determines their preferential direction of migration. The polarity vector $\vec{p}$ is reset every 1.5 minutes into a new direction using the chemokine distribution $c$ as

$$\vec{p} \;=\; \vec{p}_{\rm rand} + \frac{\alpha}{1 + \exp\left\{\kappa\left(K_{1/2} - \Delta x |\vec{\nabla} c|\right)\right\}} \frac{\vec{\nabla} c}{|\vec{\nabla} c|} \quad , \qquad \text{(S12)}$$

where $\vec{p}_{\rm rand}$ is a random polarity vector and the turning angle is sampled from the measured turning angle distribution ( (Allen *et al.*, 2007) Fig. S1B). $\alpha = 10$ determines the relative weight of the chemotaxis and random walk, $K_{1/2} = 2 \cdot 10^{11}$ Mol determines the gradient of half maximum chemotaxis weight, and $\kappa = 10^{10}/{\rm Mol}$ determines the steepness of the weight increase.

B cells de- and re-sensitize for their respective chemokine depending on the local chemokine concentration: The desensitization threshold is set to 6nMol and 0.08nMol for CXCL12 and CXCL13, respectively, which avoids cell clustering in the center of the zones. The resensitization threshold is set at $2/3$ and $3/4$ of the desensitization threshold for CXCL12 and CXCL13, respectively.

B cells can only migrate if the target node is free. If occupied and the neighbor cell is to migrate in the opposite direction (negative scalar product of the polarity vectors) both cells are exchanged with a probability of 0.5. This exchange algorithm avoids lattice artifacts leading to cell clusters.

Tfh do random walk with a preferential directionality to the LZ: The polarity vector $\vec{p}$ of Tfh is determined from a mixture of random walk $\vec{r}$ and the direction of the LZ $\vec{n}$ by

$$\vec{p} \;=\; (1 - \alpha')\vec{r} + \alpha' \vec{n}\,, \qquad \text{(S13)}$$

where $\alpha' = 0.1$ is the weight of chemotaxis. This weight leads to a dominance of random walk with a tendency to accumulate in the LZ as found in experiment. TCs migrate with an average speed of $10\mu m/min$ and repolarize every $1.7$ minutes (Miller *et al.*, 2002).

Output cell motility is derived from plasma cell motility data to be $3\mu m/min$ (Allen *et al.*, 2007) with a persistence time of $0.75$ minutes.

#### Mutation distance Shannon Entropy and shape space point Shannon Entropy calculation

In addition to fcSE we also calculate two other diversity measures namely, mutation distance Shannon Entropy and shape space point Shannon Entropy (SSpoint SE). These two diversity measures are calculated similar to Shannon Entropy. At time t:

$$\mathrm{SE} = -\sum_{i=1}^{N} p_\mathrm{i} * \log_\mathrm{e}(p_\mathrm{i})$$

and $p_\mathrm{i}$ is defined as:

$$p_\mathrm{i} = \frac{\chi_\mathrm{i}}{\chi_\mathrm{total}}$$

For mutation distance Shannon Entropy, N is the total number of unique mutation distances of B cells and $\chi_\mathrm{i}$ is the number of B cells having $i^\mathrm{th}$ mutation distance at time t. For Shape Space point Shannon Entropy, N is the total number of unique points on the shape space occupied by B cells and $\chi_\mathrm{i}$ is the number of B cells occupying $i^\mathrm{th}$ point in the shape space at time t. $\chi_\mathrm{total}$ is the total number of B cells at time t. As an example, analysis for antigen availability using these two diversity measures are reported in (Fig. S1C and D) .

### 2 SUPPLEMENTARY TABLES AND FIGURES

#### 2.1 Table: Systems parameters

| Index | Name | Default value | Results section |
|---|---|---|---|
| 1 | Initial antigen amount on FDC | 3500 units | 2.2 – 2.5 |
| 2 | Mutation distance of founder cells | 5 to 6 | 2.1, 2.3 – 2.5 |
| 3 | Number of Tfh cells | 200 | 2.1 – 2.3, 2.5 |
| 4 | Tfh signal threshold | 0.5 hrs | 2.1 – 2.5 |
| 5 | Tfh signal multiplier | 1 | 2.1 – 2.4 |
| 6 | DND function: K value | 1.3 hrs | 2.1 – 2.4 |
| 7 | DND function: Hill coeff. | 1.8 | 2.1 – 2.4 |

**Table S1.** Table of key simulation parameters.

#### 2.2 Supporting Figures

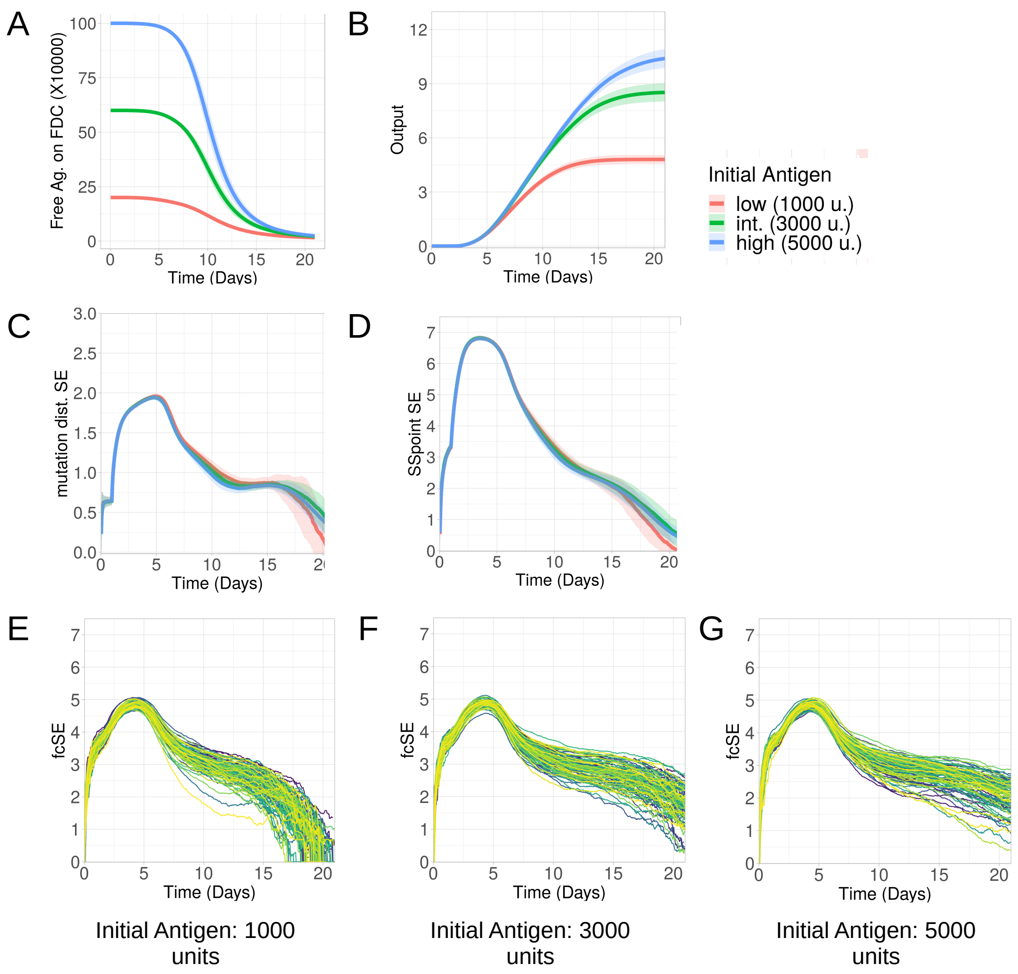


**Figure S1.** Simulation with changing antigen amounts related to (see the main text, Fig. 1). (A) Free Ag on FDC, (B) GC output, (C) mutation distance Shannon Entropy, (D) Shape Space point Shannon Entropy, (E), (F) and (G) founder cell Shannon Entropy (fcSE) for each individual GC for low, intermediate and high antigen. Mean (continuous lines) and standard deviation (shaded area) of simulations for a total of 20 simulated GCs are shown.

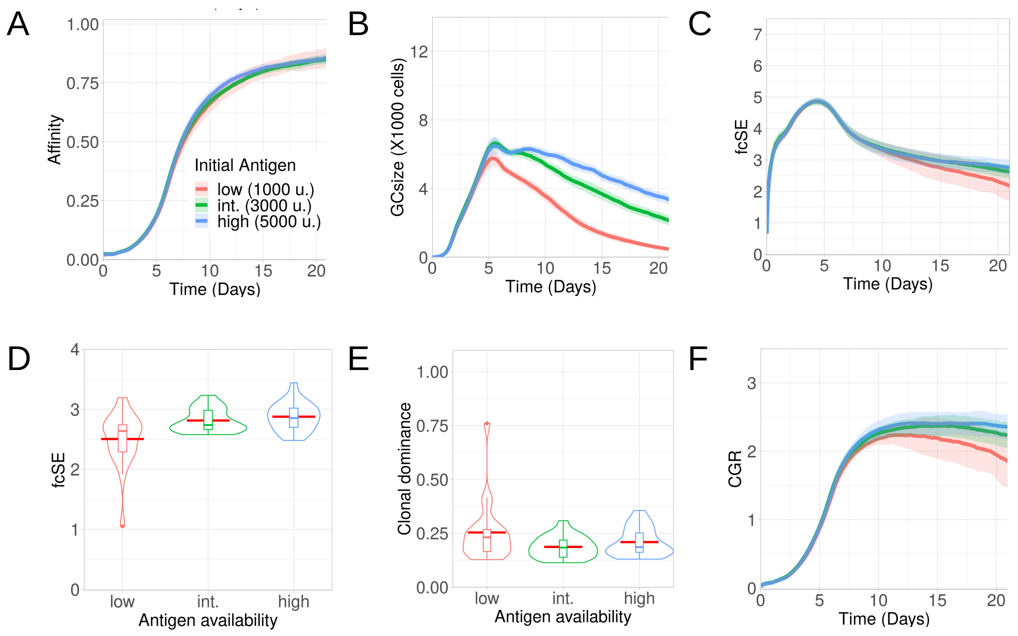


**Figure S2.** Simulations done without having endogenous antibody feedback and changing antigen amounts as low (1000 units in red), intermediate (3000 units in green) and high (5000 units in blue): (A) Affinity of GC B cells, (B) GC size, (C) founder cell Shannon Entropy (fcSE), (D) violin plot of fcSE at day 18, (E) violin plot of clonal dominance at day 18 and (F) cumulative GC response (CGR). Mean (continuous lines) and standard deviation (shaded area) of simulations for a total of 20 simulated GCs are shown. The box plots of the violin plots show the median (horizontal line inside the box), 25 and 75 percentiles, the mean (horizontal red line) and the outlier points as dots.

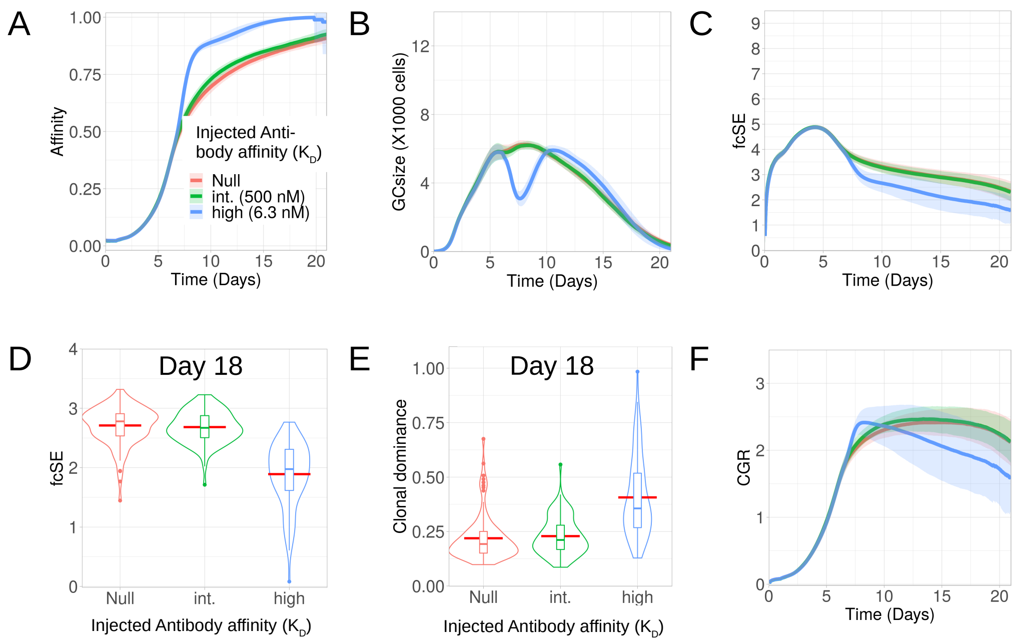


**Figure S3.** Simulations done with changing affinity of exogenous antibody (antibody injection at day 6) as Null (no injection in red), intermediate (500 nM in green) and high (6.3 nM in blue): (A) Affinity of GC B cells, (B) GC size, (C) founder cell Shannon Entropy (fcSE), (D) violin plot of fcSE at day 18, (E) violin plot of clonal dominance at day 18 and (F) cumulative GC response (CGR). Mean (continuous lines) and standard deviation (shaded area) of simulations for a total of 100 simulated GCs are shown. The box plots of the violin plots show the median (horizontal line inside the box), 25 and 75 percentiles, the mean (horizontal red line) and the outlier points as dots.

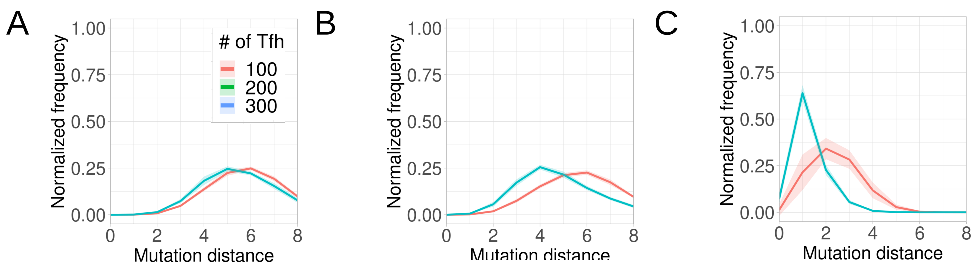


**Figure S4.** Simulations done with changing Tfh numbers. Normalized frequencies of GC B cells of different mutation distances at (A)3 days (B)4 days and (C)10 days

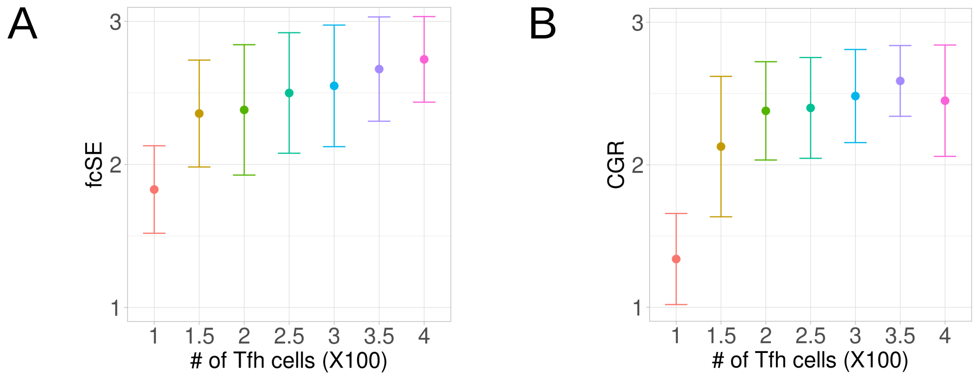


**Figure S5.** Simulations done with changing Tfh numbers. (A) founder cell Shannon Entropy (fcSE) at day 18 (B) cumulative GC response (CGR) at day 18.

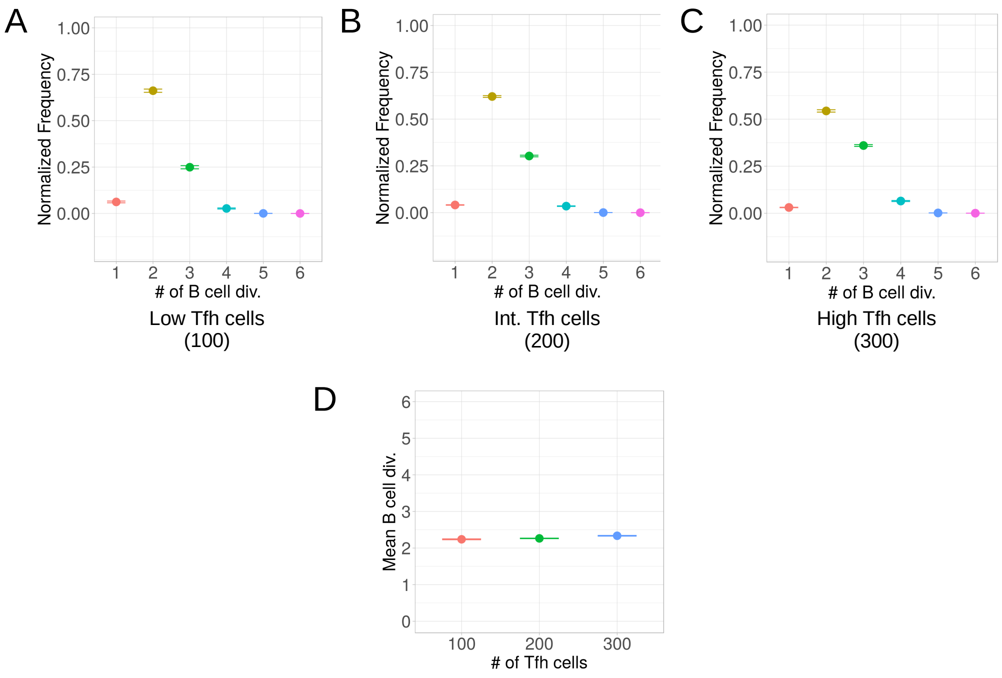


**Figure S6.** Simulations done with changing Tfh cell numbers. Normalized frequencies of number of divisions of B cells for (A)100 Tfh (B)200 Tfh and (C)300 Tfh. (D) number of mean B cell divisions

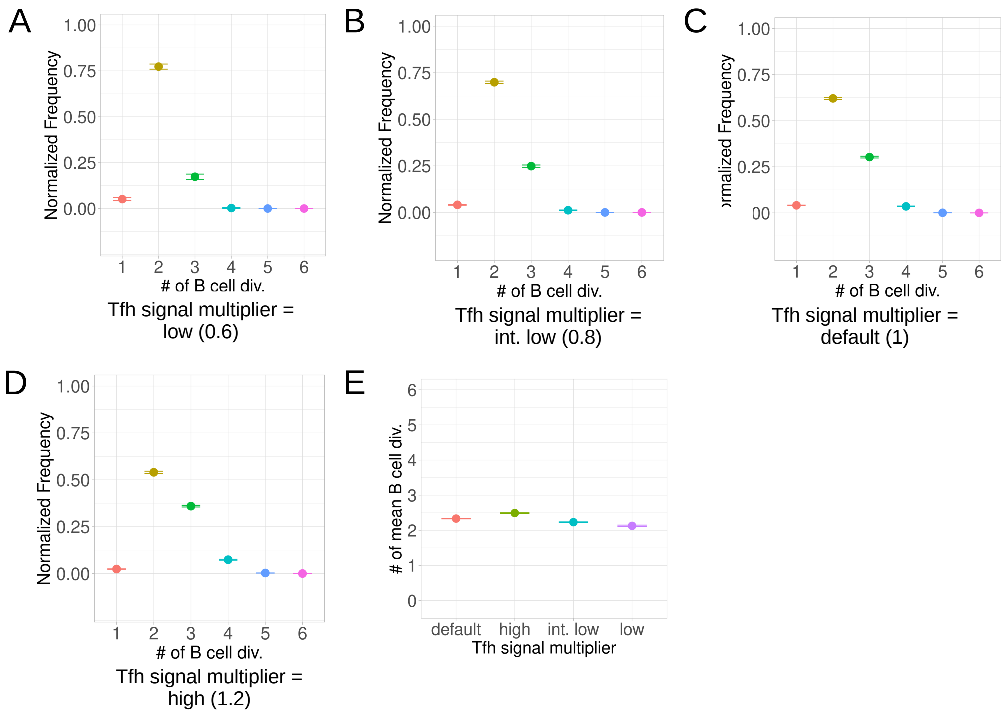


**Figure S7.** Simulations done with changing Tfh signal multiplier. Normalized frequencies of number of divisions of B cells for Tfh signal multiplier value of: (A) low (0.6), (B) intermediate low (0.8), (C) default (1.0) and (D) high (1.2). (E) number of mean B cell divisions
